# Cerebral ischemia induces TRPC6 in the glomerular podocytes: A novel role for HIF1α/ZEB2 axis in the pathogenesis of stroke-induced proteinuria

**DOI:** 10.1101/538959

**Authors:** Krishnamurthy Nakuluri, Rajkishor Nishad, Dhanunjay Mukhi, Sireesh Kumar, Venkata P Nakka, Lakshmi P Kolligundla, Parimala Narne, Sai Sampath K Natuva, Prakash Babu Phanithi, Anil K Pasupulati

## Abstract

Glomerular filtration apparatus (GFA) regulates the glomerular permselectivity and ultrafiltration of urine. Podocytes are specialized cells and a key component of the GFA. The mechanism by which the integrity of the GFA is compromised and manifest in proteinuria during ischemic stroke remains enigmatic. Hypoxia is a determining factor in the pathophysiology of ischemia. We investigated the mechanism of ischemic-hypoxia induced proteinuria in a middle cerebral artery occlusion (MCAO) model. Ischemic hypoxia resulted in the accumulation of HIF1α in the glomerular podocytes that resulted in the increased expression of ZEB2. ZEB2, in turn, induced TRPC6 (transient receptor potential cation channel, subfamily C, member 6), which has increased selectivity for calcium. Elevated expression of TRPC6 elicited increased calcium influx and aberrant activation of focal adhesion kinase (FAK) in podocytes. FAK activation resulted in the stress fibers reorganization and podocyte foot process effacement. Our study suggests overactive HIF1α/ZEB2 axis during ischemic-hypoxia induces intracellular calcium levels via TRPC6 and consequently altered podocyte integrity and permselectivity.

## Introduction

Extreme physiological and pathological conditions impose challenges on human physiology. The normal functioning of the human body demands both continuous and adequate supply of oxygen whereas relative (hypoxia) and the absolute deficiency (anoxia) of oxygen are a risk to human health. Human organs vary in their oxygen dependency and susceptibility to oxygen deficiency. Brain and kidney are most hypoxia-sensitive organs. Formation of ATP from ADP is a major chemical reaction where oxygen is involved. ATP-dependent active salt re-absorption in kidney demands high oxygen supply^1^. Kidney carries out its functions within a narrow range of partial pressure of oxygen, which is very low in the inner medulla (5 mmHg) and compared with the outer cortex (50 mmHg)^2^. Furthermore, renal vasculature despite its low-resistance subjected to continuous perfusion with a high-volume^3,4^. Both, vascular architecture of the kidney and excess demand for oxygen integrate and let the kidneys become more sensitive to oxygen-deprived conditions^1,5,6^. Limitations in oxygen supply impose kidneys to undergo hypoxia-induced maladaptation, which likely reflects in the pathophysiology of acute kidney injury and proteinuria^6–12^.

The vertebrate kidneys regulate homeostasis predominantly by controlling acid-base, electrolyte, and water balance. Kidneys are also instrumental in ultrafiltration of plasma components and regulating the composition of urine. Proteinuric condition suggests abnormalities in the glomerular filtration apparatus (GFA)^13^. Three layers of GFA are podocytes, glomerular basement membrane and, perforated endothelium^13^. Proteinuria is often associated with clinical conditions such as stroke and sleep apnea. These conditions are presented with reduced renal perfusion and moderate to severe hypoxia^12,14^. Accumulated evidence suggests that hypoxia contributes to the proteinuria and pathogenesis of chronic kidney disease (CKD)^6,7,10,15–17^. The prevalence of CKD is more than 30% among stroke patients^18^. Renal dysfunction is a worse clinical outcome in patients with ischemic stroke^19,20^ and it is also an independent predictor of stroke mortality^18^.

Epidemiological studies revealed that almost 1 in 10 adults in the USA has proteinuria^21^. Owing to the large volume of proteinuric population and the incidence of proteinuria with the disorders wherein anoxia/hypoxia prevail, it is crucial to gain insights on the pathophysiological relation between proteinuria and hypoxia. In this study, we employed middle cerebral artery occlusion (MCAO) model, which is routinely employed to mimic human ischemic stroke and investigated the mechanism of glomerular dysfunction. We found that ischemic stroke and the resultant ischemic-hypoxia injury resulted in the elevated expression of HIF1α in various organs including kidney. ZEB2, a downstream target of HIF1α is induced in glomerular podocytes, which in turn induced the expression of TRPC6 and calcium influx in podocytes. Calcium-induced FAK activation resulted in stress fibers rearrangement and manifested in impaired podocyte structure and function. Overactivity of the HIF1α/ ZEB2 axis and TRPC6 could be a mechanism by which systemic hypoxia manifests in proteinuria.

## Results

### Ischemic stroke is associated with systemic hypoxia and proteinuria

We performed triphenyl tetrazolium chloride (TTC) staining to assess the infarct size and injury following MCAO and reperfusion. Ischemic region of the brain, when stained with TTC, showed white color indicating the infarct lesions (Fig 1A). 24 h following the MCAO surgery in rats, we measured the urine albumin to creatinine ratio (ACR) to evaluate the effect of ischemic stroke injury on renal function. The ischemic stroke resulted in significant albuminuria compared with sham-operated rats (Fig.1B). Silver staining revealed a significant amount of protein (particularly albumin) in urinary fractions from stroke-induced rats compared with sham-operated rats (Fig.1C). Further, to ascertain whether ischemic-reperfusion injury elicits hypoxia, we analyzed the expression of HIF1α in brain and kidney and found that HIF1α is elevated in these organs (Fig.1D-G). The data suggest that in ischemic stroke rats elevated renal expression of HIF1α is concurrent with the adverse renal outcome.

**Figure 1.**
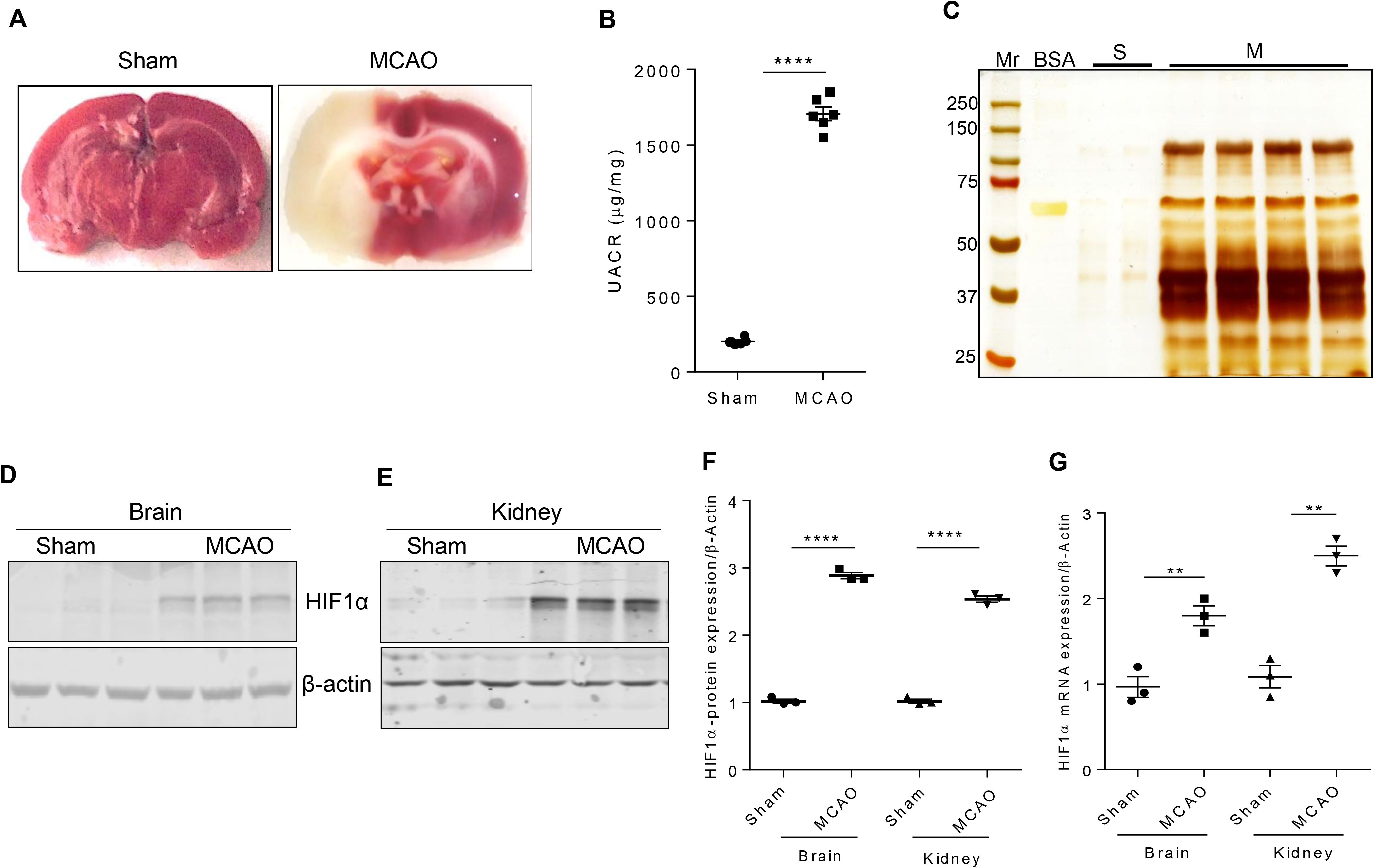
Ischemic stroke alters kidney function. (A) TTC staining images of sham and ischemic stroke-induced (MCAO) rat brain. (B) Estimation of albumin and creatinine levels in MCAO rats. Error bars indicate mean ± SE; n=6. ****p<0.001. (C) Urine samples from sham (S) and stroke-induced rats (M) were subjected to SDS-PAGE and band were visualized by silver staining, Mr, molecular weight marker (#1610374; Bio-Rad); BSA-Bovine Serum Albumin. HIF1α expression in the infarcted region of the brain (D) and glomerular lysates from sham and MCAO rats (E). Densitometric analysis of HIF1α band is depicted after normalized for respective β-actin expression (F). Error bars indicate mean ± SE; n=3. ****p<0.001. (G) Steady-state mRNA levels of HIF1α from the brain and glomerular lysates were measured by qRT-PCR. Error bars indicate mean ± SE; n=3. **p<0.003.

Since we observed proteinuria in stroke-induced rats we performed various staining procedures to assess stroke-induced renal morphological changes. PAS staining did not reveal glomerulosclerosis in stroke-induced rats. Owing to the importance of podocytes in glomerular filtration we assessed the podocyte number and their morphology in experimental rats. WT1 staining revealed a decreased number of podocytes in stroke-induced rats (Fig.2A). Transmission electron microscope images revealed shortened podocyte foot-processes and an increase in the thickness of the basement membrane (Fig.2B).

**Figure 2.**
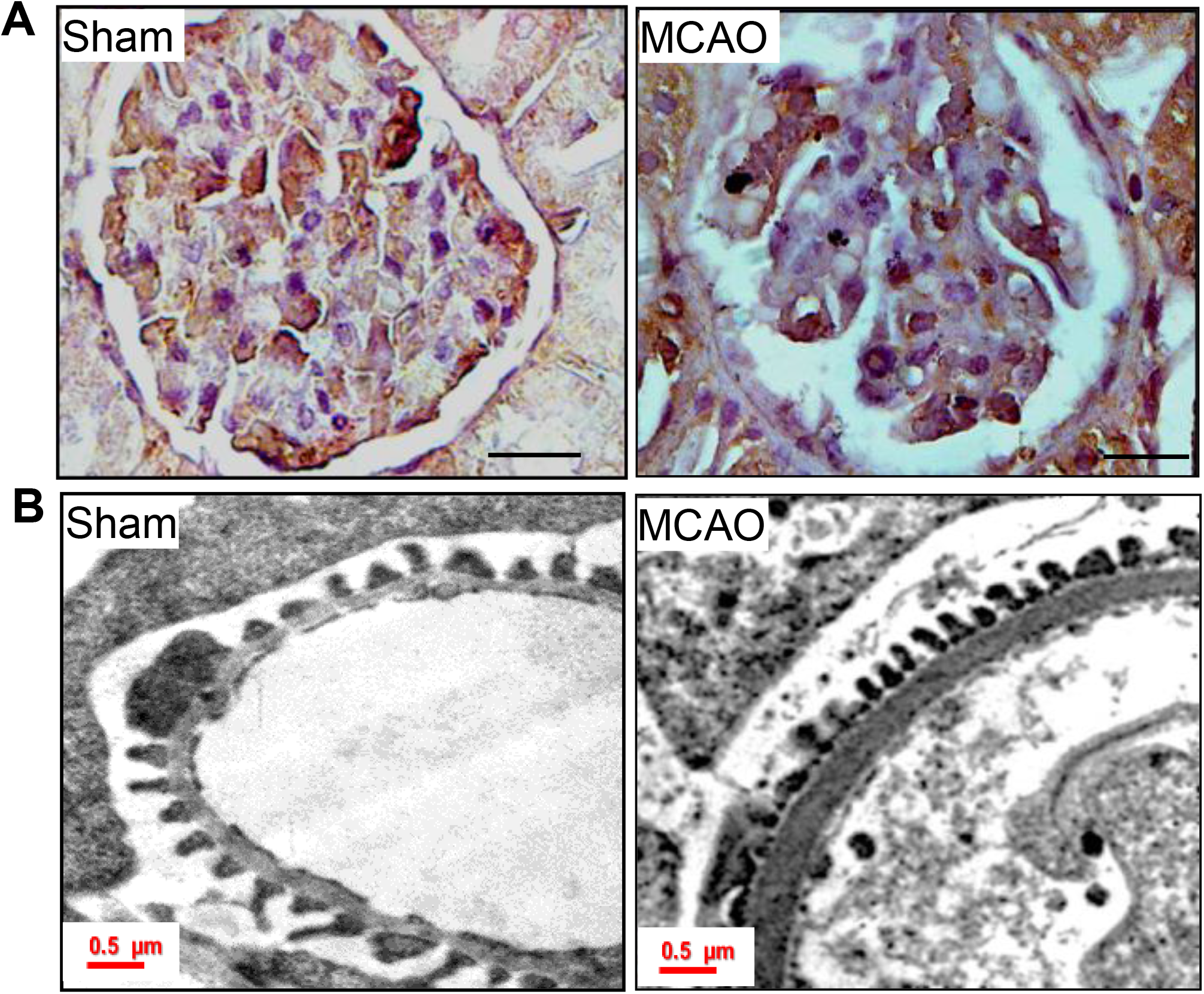
Ischemic-hypoxia elicits podocyte injury. (A) Staining for WT1 in sham and MCAO rat glomerular sections. Images were obtained with a 100x objective Leica trinocular microscope. (B) TEM images of podocyte foot processes in sham and MCAO rat kidney sections. In ischemic-stroke rats, podocyte foot-processes were small and the thickness of GBM was increased compared to sham-operated rats.

### Ischemic hypoxia induces ZEB2 and TRPC6 expression in podocytes

It was shown earlier that elevated ZEB2 expression is concomitant with HIF1α accumulation in podocytes exposed to hypoxia^23^. ZEB2, a zinc-finger transcription factor regulates the expression of several proteins representing various cellular processes including cell adhesion and epithelial-mesenchymal transition (EMT). Fittingly, in our stroke-induced rats, we observed that ZEB2 was induced along with HIF1α in the glomerular podocytes (Fig.3A, C&E). Increased expression of HIF1α and its down-stream target ZEB2 is associated with reduced expression of E-cadherin in podocytes isolated from MCAO rats and human podocytes treated with FG-4592 (Fig.3A-D). FG-4592 is a prolyl hydroxylase inhibitor, which stabilizes the HIF1α expression and elicits activation of target genes and signaling events. E-cadherin is a bona fide target of ZEB2 and attenuation of E-cadherin is a characteristic feature of EMT^24–26^. Loss of ZEB2 in epithelial cells showed migration defects, which was associated with decreased expression of TRPC6 (transient receptor potential cation channel, subfamily C, member 6)^27^. Promoter analysis (TFSearch) showed that ZEB2 occupies E2-box1 (−430CAGGTG-425) of the of human TRPC6 promoter. Further, we also found that TRPC6 expression was elevated in both stroke-induced rat podocytes and human podocytes exposed to FG-4592 (Fig.3A-D). Immunostaining data revealed that HIF1α, ZEB2, and TRPC6 were induced in glomeruli from stroke-induced rats (Fig.3E). Immunofluorescence data suggests co-localization of HIF1α and ZEB2 in podocytes treated with FG-4592 (Fig.3F). Further, we also demonstrated elevated expression of ZEB2 and TRPC6 in FG-4592 treated podocytes (Fig.3G). Together, the data suggest TRPC6 expression is concomitant with ZEB2 expression in podocytes with elevated HIF1α expression.

**Figure 3.**
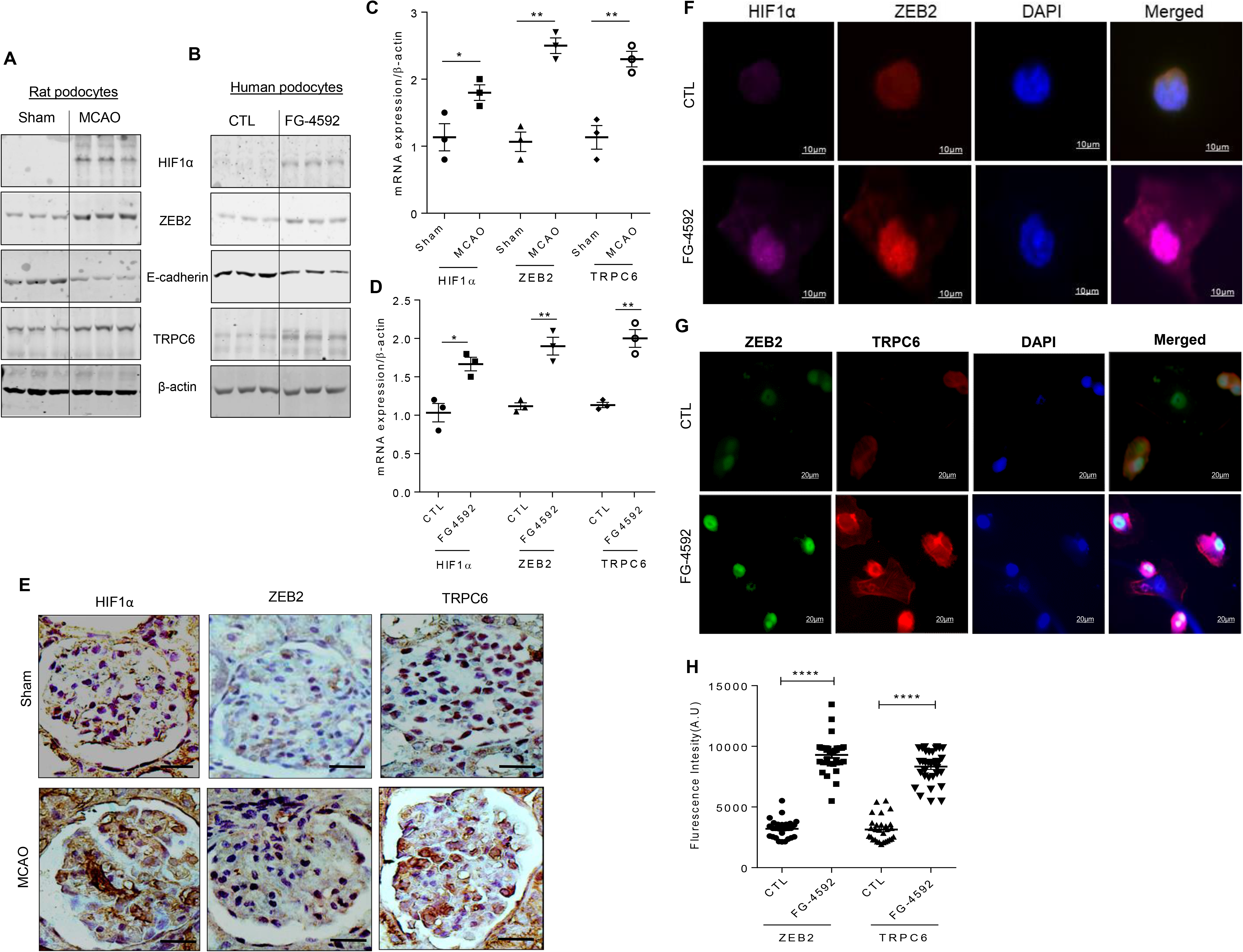
Ischemic-hypoxia induces ZEB2 and its target genes in glomerular podocytes: (A) Lysate from primary podocyte isolated from sham and MCAO rat kidney was used to assess the expression of HIF1α, ZEB2, E-cadherin, and TRPC6. (B) Differentiated human podocytes treated with or without FG-4592 and analyzed the expression of HIF1α, ZEB2, E-cadherin and TRPC6. mRNA levels of HIF1α, ZEB2, and TRPC6 from podocytes isolated from sham and MCAO rats (C) and human podocytes treated with or without FG-4592 (D). Error bars indicate mean ± SE; n=3. *p<0.05; **p<0.003. (E) Immunohistochemical analysis of HIF1α, ZEB2, and TRPC6 in glomerular sections from sham and MCAO rats. The scale bar represents images of 20μm and images were captured with a 100x objective of Leica trinocular microscope. (F) Co-localization of HIF1α and ZEB2 in human podocytes treated with or without FG-4592. Images were acquired using a Zeiss 100x objective. (G) Elevated expression of ZEB2 and TRPC6 in human podocytes treated with FG-4592. Images were acquired with a Leica trinocular 63x objective and (H) fluorescence intensity of ZEB2 and TRPC6 expression in podocytes treated with or without FG-4592. Error bars indicate mean ± SE; n=20. ****p<0.001.

### ZEB2 induces TRPC6 expression and calcium influx in podocytes

We performed ChIP assay to demonstrate the interaction between TRPC6 promoter and ZEB2. We found that ZEB2 binds to the proximal promoter of TRPC6 at the E2-box1 region during hypoxic conditions (Fig.4A). We used E-cadherin as a positive control based on earlier studies that reported E-cadherin is suppressed by ZEB2^24^. We also confirmed HIF1α occupancy on ZEB2 promoter (Fig.4A) wherein, the interaction between HIF1α with VEGF promoter serves as a positive control. Furthermore, we observed increased promoter activity of TRPC6 in HEK293T cells treated with either FG-4592 (Fig.4B) or with ectopic expression of ZEB2.

Upon confirming that accumulation of HIF1α and ZEB2 is followed by TRPC6 expression, we investigated the significance of elevated expression of TRPC6 in podocytes. Although TRP family proteins have an affinity for cation transport, TRPC6 has selectivity for calcium influx. Among glomerular cells, TRPC6 expresses predominantly in podocytes^28^. We found increased intracellular calcium influx in podocytes exposed to FG-4592 as measured by calcium-sensitive fluorescent dye Fluo3-AM (Fig.4C). While ectopic expression of ZEB2 increased calcium influx, either knockdown of ZEB2 or exposure of cells to calcium channel blocker 2-aminoethoxy diphenylborate (2APB) resulted in decreased intracellular calcium levels in podocytes (Fig.4C).

**Figure 4.**
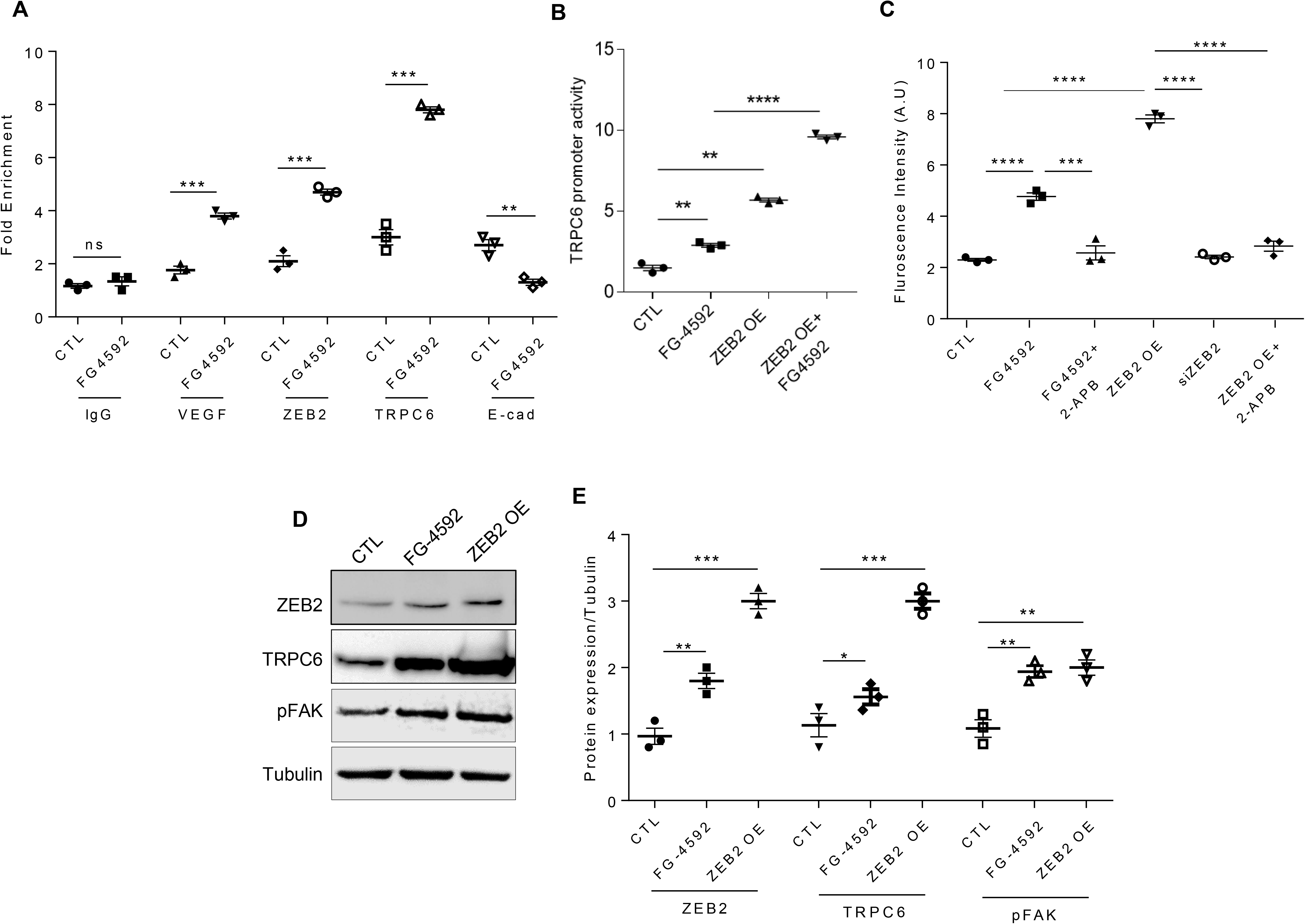
Hypoxia induces HIF1α-ZEB2-TRPC6 axis: (A) ChIP analysis with chromatin fractions from podocytes exposed to FG-4592 was performed as described in methods. Input DNA and DNA from each of the immunoprecipitated samples were PCR amplified for hypoxia response element (HRE) in VEGF promoter and ZEB2 promoter and E2-box region in both TRPC6 promoter and E-cadherin promoter. HRE in VEGF promoter fragment and E2-box in E-cadherin promoter serve as positive controls for HIF1α binding and ZEB2 binding respectively. Error bars indicate mean ± SE; n=3. **p<0.0 03; ***p<0.02. (B) TRPC6 promoter activity was measured in HEK293T cells that ectopically expressing ZEB2 and exposed to FG-4592. Renilla luciferase was used as an internal control to normalize transfection efficiency. Error bars indicate mean ± SE; n=3. **p<0.003; ***p<0.002. (C) Intracellular calcium levels in podocytes were measured by Fluo3-AM following treatment with FG-4592 in the presence or absence of 2-APB. Podocytes expressing siZEB2 and ectopically expressing ZEB2 were employed in this study. Error bars indicate mean ± SE; n=3. **p<0.003; ***p<0.002, ****p<0.001. (D) Immunoblot analysis of ZEB2, TRPC6, and pFAK in podocytes treated with FG-4592 or ectopically expressing ZEB2 (ZEB2 OE). (E) Quantification of band intensities of ZEB2, TRPC6, pFAK was Image J analysis (NIH). Error bars indicate mean ± SE; n=3. *p<0.05, **p<0.003, ***p<0.002, ****p<0.001.

### ZEB2 regulates activation of FAK via TRPC6

Elevated intracellular calcium levels elicit auto-phosphorylation (Y397) of focal adhesion kinase (FAK)^29^. On the other hand, inhibition of FAK protects against effacement of podocyte foot-processes^30^. FAK is a central protein of focal adhesions and is known to regulate the function of cytoskeletal and focal adhesion proteins^31^. Therefore, we measured pFAK levels in podocytes ectopically expressing ZEB2 or treated with FG-4592. ZEB2 overexpression in podocytes resulted in both increased TRPC6 expression and activation of FAK (Fig 4D&E). On the other hand, ZEB2 knockdown is associated with reduced expression of TRPC6 and pFAK (Fig.5A&B). Together, the data suggest that ZEB2 regulates phosphorylation of FAK via TRPC6. Further, to ascertain the essential role of TRPC6 in FAK activation, we attenuated TRPC6 expression by siTRPC6 and measured the pFAK levels. pFAK levels were decreased in relation to TRPC6 levels in cells treated with or without FG-4592 (Fig 5C, D, S1&2).

**Figure 5.**
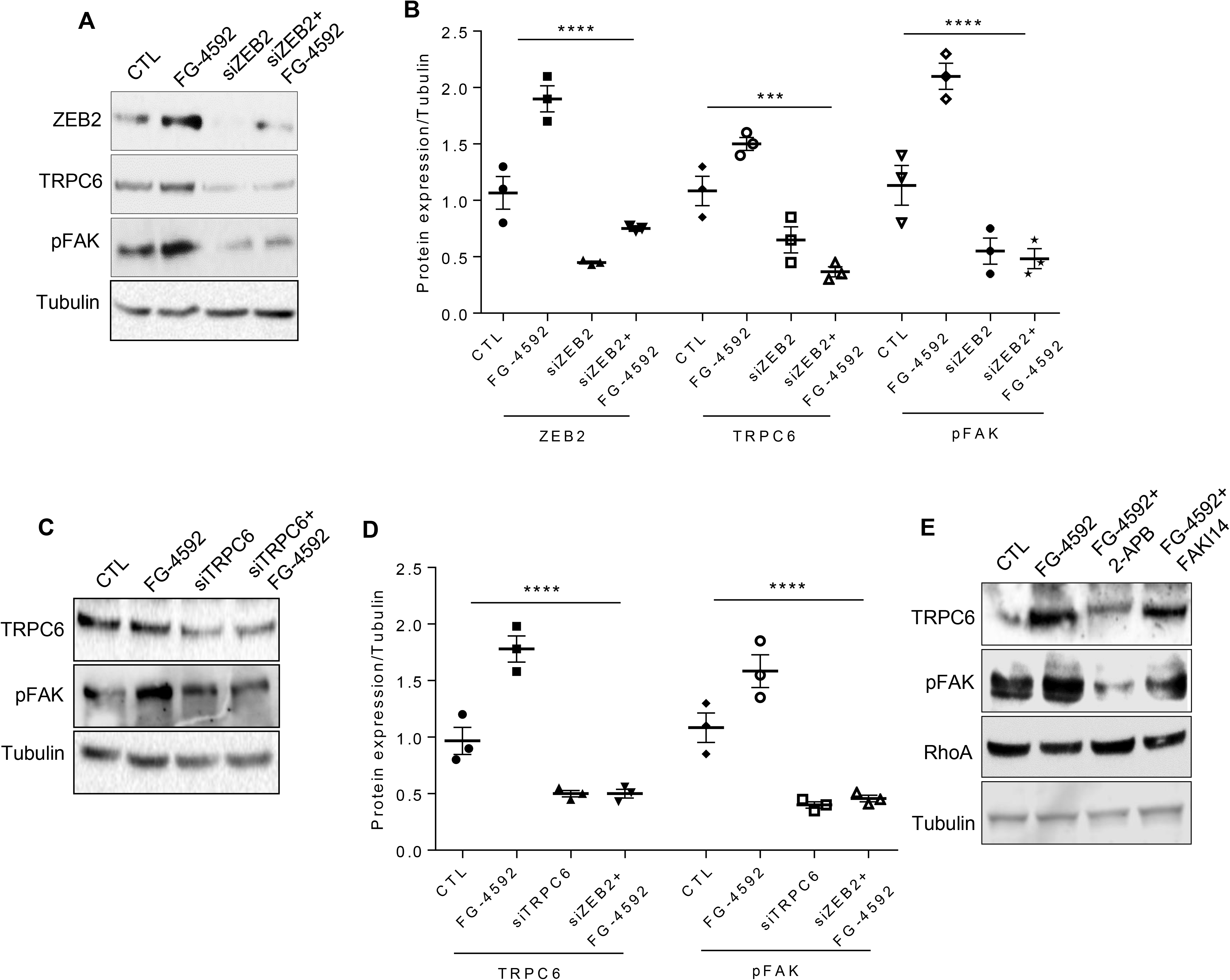
Essential role of ZEB2 in regulating TRPC6 expression: (A) Immunoblotting analysis of ZEB2, TRPC6, and pFAK expression in podocytes expressing siZEB2 and treated with or without FG-4592. (B) Quantification of band intensities of ZEB2, TRPC6, pFAK was performed with Image J software. Error bars indicate mean ± SE; n=3, ****p<0.00 1. (C) Immunoblotting analysis of TRPC6 and pFAK expression in podocytes in which TRPC6 expression was knocked-down and treated with or without FG-4592 (D) Quantification of band intensities of western blots was performed with Image J. Error bars indicate mean ± SE; n=3, ****p<0.001. (E) Immunoblotting analysis of TRPC6, pFAK, and RhoA expression in podocytes exposed to FG-4592 and treated with or without 2APB and FAKI14.

Activated FAK is restricted to focal adhesions. Focal adhesions serve as connections between the cytoskeleton and the cell matrix, whereas their formation is regulated by RhoA^31^. FAK suppresses RhoA activity and promotes focal adhesion turnover^31,32^. In similar to the earlier reports, we also noticed decreased RhoA while FAK gets activated in podocytes treated with FG-4592 (Fig 5E). RhoA expression was restored in podocytes treated with either calcium channel blocker (2APB) or FAK inhibitor14 (FAKI14) (Fig 5E).

### HIF1α alters podocyte actin cytoskeleton

As we noticed activation of FAK and decreased RhoA expression in podocytes with elevated HIF1α, we next investigated for the cytoskeletal abnormalities, if any. We assessed the actin stress fibers (SFs) distribution in cells that are naïve or exposed FG-4592 employing Phalloidin staining. Differentiated podocytes are naïve to hypoxia exhibit orderly arranged non-branching SFs (Fig. 6Ai). Upon exposure to hypoxia, the orderly arranged stress fibers of the podocyte actin cytoskeleton were disrupted (Fig. 6Aii). Stress fibers reorganization and altered morphology was also observed in cells that ectopically express ZEB2 and treated with or without FG-4592 (Fig.6Aiii&iv). Interestingly, ZEB2 knockdown improved reorganization of SFs in podocytes exposed to FG-4592 (Fig 6Av). In podocytes that were exposed to FG-4592, or ectopically expressing ZEB2, treatment with 2APB or FAKI14 improved reorganization of SFs (Fig.6Avi-ix). Podocytes in which TRPC6 was knocked down, FG-4592 treatment did not alter the distribution of SFs (Fig.6Ax). We quantified the number of SFs per podocyte and the ratio of SFs to total cell size was calculated using Image J (NIH). The data (Fig.6B) suggests that inhibition of TRPC6 channel thus calcium influx or inhibition of FAK could prevent HIF1α induced podocyte cytoskeletal rearrangements. The disruption of SFs of the podocyte actin cytoskeleton may suggest the reason for altered cell morphology and FPE in podocytes from stroke-induced rats (Fig 2B vs. 6Aii-iv).

**Figure 6.**
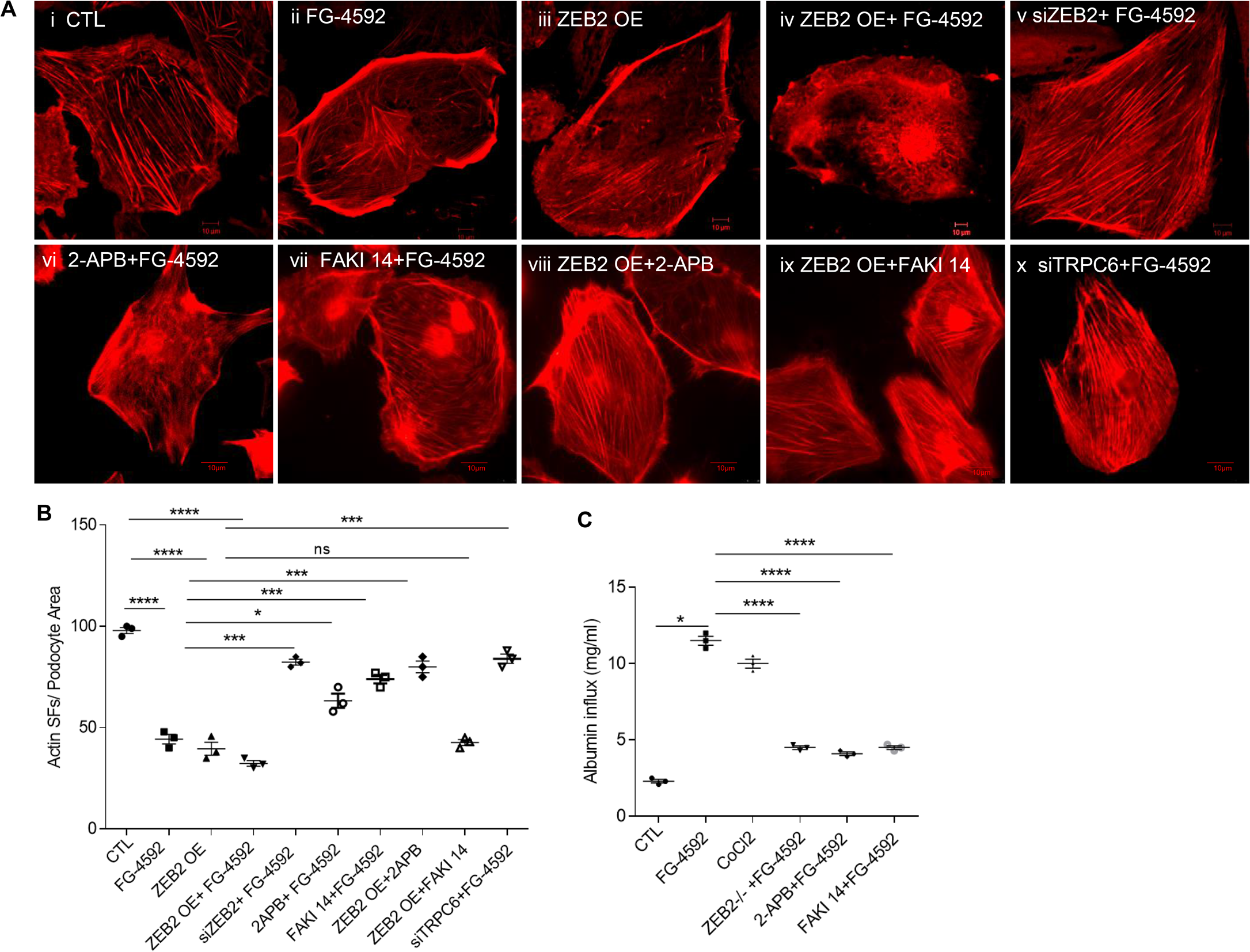
HIF1α-ZEB2-TRPC6 axis regulates podocyte cytoskeleton reorganization: (A) Phalloidin staining was performed to visualize intracellular F-actin stress fibers (SFs) in podocytes. Podocytes treated with FG-4592 (ii) compromised their morphology and SFs arrangement compared with cells naïve to FG-4592 (i). Ectopic expression of ZEB2 with and without FG-4592 treatment also resulted in altered SFs arrangement (iii & iv). Knockdown of ZEB2 in podocytes by siZEB2 (v) or podocytes treated with calcium channel blocker; 2APB or FAK inhibitor; FAKI14 (vi & vii) improved SFs reorganization. Both 2APB and FAKI14 ameliorated SFs organization in podocytes, which overexpressing ZEB2 (viii&ix). Knockdown of TRPC6 expression in podocytes by siTRPC6 also ameliorated SFs reorganization (x). The scale bar represents images of 50μm and images were captured with a 63x objective of Leica trinocular microscope. (B) The number of stress fibers per podocyte was quantified and analyzed by Image-J (NIH) and the ratio of stress fibers to total cell size was represented. Error bars indicate mean ± SE; n=3. *p<0.05. (C) Quantification of albumin influx across podocyte monolayer. Podocytes transduced with or without ZEB2 were grown as a monolayer on cell culture insert and allowed to differentiate. Differentiated podocytes were exposed to FG-4592 for 24 h and treated with FAKI14 or 2APB and albumin influx assay was performed as described in methods. Error bars indicate mean ± SE; n=3. *p<0.05. Podocytes transduced with shZEB2 or treated with FAKI14 or 2APB showed improved permselectivity to albumin than podocytes exposed to FG-4592 alone.

### Calcium channel blocker and FAK inhibitor prevent HIF1α induced podocyte permeability

Earlier studies revealed that accumulation of HIF1α in podocytes increased podocyte permeability to albumin and blunting the expression of ZEB2 diminished the susceptibility podocyte monolayer to hypoxia-driven elevated albumin influx^23^. This data strongly suggest that ZEB2 is necessary to elicit HIF1α’s action on podocyte permeability. Since we showed that ZEB2 induces TRPC6 and calcium accumulation and consequent activation of FAK in podocytes, we assessed the permeability of podocytes treated with FG-4592 in the absence or presence of 2APB and FAKI14. Both 2APB and FAKI14 prevented the podocyte permeability to albumin (Fig.6C), suggesting that prevention of calcium influx and/or inhibition of FAK preserve podocyte permselectivity during hypoxia.

### Co-expression of HIF1α, ZEB2, and TRPC6 in kidney diseases

It is noteworthy that intrarenal hypoxic injury is considered to be the common cause of renal dysfunction and proteinuria in conditions such as hypertension, diabetes, and stroke^33,34^. We performed co-expression analysis for HIF1α, ZEB2, and TRPC6 in the *Nephroseq* database (University of Michigan, Ann Arbor). *Nephroseq* analysis revealed increased expression of HIF1α, ZEB2, and TRPC6 in Nakagawa CKD dataset and in Hodgin Diabetes Mouse Glomeruli datasets (Fig.7A&B). The data suggests these three genes co-express in CKD of human origin and in diabetic mouse glomerular diseases.

**Figure 7.**
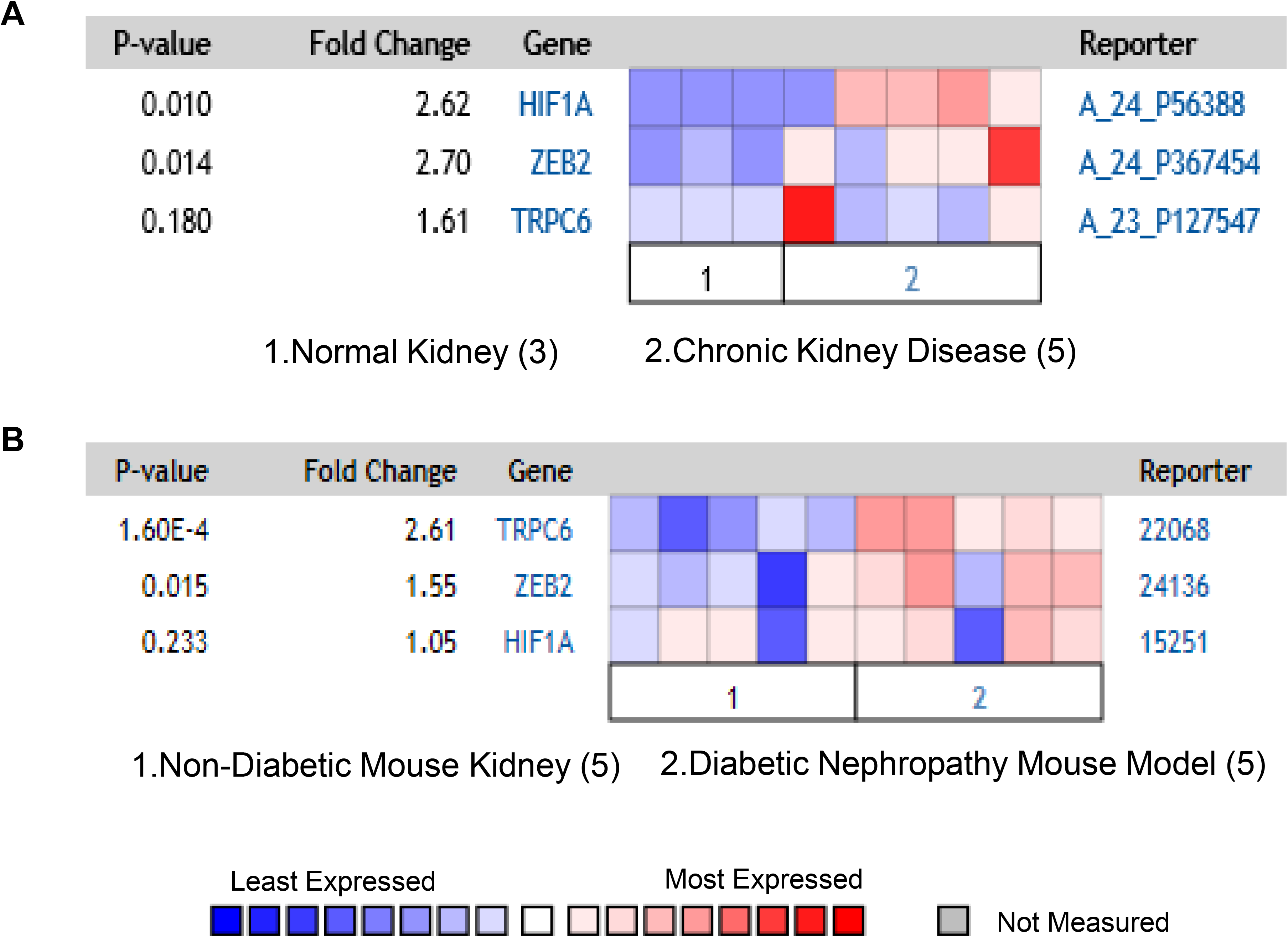
Co-expression of HIF1α, ZEB2, and TRPC6 in glomerular diseases. (A) Nakagawa CKD data set showing the elevated expression of HIF1α (2.6 fold), ZEB2 (2.7 fold), and TRPC6 (1.6 fold) in patients with chronic kidney disease vs. healthy kidney. (B) Hodgin diabetes mouse glomeruli datasets showing the elevated expression of ZEB2 (1.55 fold), and TRPC6 (2.61 fold) in mouse with diabetic nephropathy vs. non-diabetic mouse models. The data is obtained from *Nephroseq* (University of Michigan, Ann Arbor, MI).

### Calcium channel blockers improved proteinuria in ischemic stroke patients

120 ischemic stroke patients with and without hypertension were admitted in this study and those patients who had the cerebrovascular disease, cardiovascular disease, and diabetes were excluded (Table S1&2). Ischemic stroke patients without hypertension history (n=50) were followed for a mean period of three years. Previous studies have shown that ischemic stroke is accompanied by proteinuria and reduced renal outcome^35,36^. We collected the data from ischemic stroke patients before and after treating with calcium channel blockers. Ischemic stroke patients have shown reduced eGFR (Fig.8A, <90), increased albuminuria, and proteinuria (Fig.8B&C). Treating stroke patients with dihydropyridine, an analog of calcium channel blocker with other supplements of therapeutic regimen improved renal function in proteinuric patients (Fig.8&Table S2). This data supports our animal experiment where we observed ischemic reperfusion injury is associated with proteinuria and calcium blockers may improve renal outcome.

**Figure 8:**
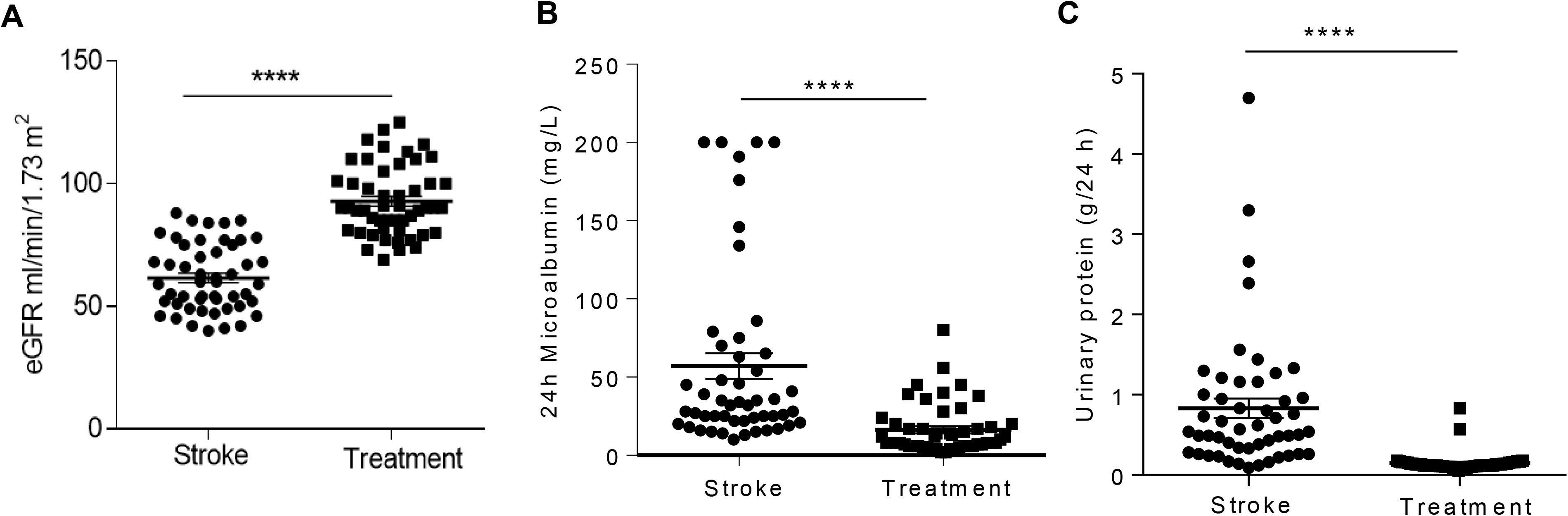
The therapeutic regimen with calcium channel blockers improved proteinuria in ischemic-stroke patients: (A) eGFR (B) 24h urinary microalbumin (mg/L) and (C) 24h urinary protein concentrations were measured in ischemic stroke patients before and after treatment with regimen composed of dihydropyridines, a calcium channel blockers. Error bars indicate mean ± SE; n=50. ****p<0.0 01.

## Discussion

Podocytes are instrumental for contributing glomerular permselectivity and ultrafiltration of urine. It has been known that ischemic stroke is often associated with proteinuria. Owing to the importance of podocytes in glomerular filtration, we investigated the cellular effects of stroke-associated ischemia-hypoxia on podocyte biology. We show that following ischemic reperfusion, HIF1α and its down-stream target ZEB2 are elevated in glomerular region and especially in podocytes. Our results suggest a novel role of HIF1α with the elevated expression of TRPC6 in podocytes. Elevated expression of TRPC6 is at least partially due to ZEB2 expression. TRPC6 ensures calcium influx into podocytes, which elicits FAK activation and these events culminate in the disruption of actin stress fibers. In addition to altered morphology of podocytes, accumulation of HIF1α resulted in the increased permeability to albumin across podocyte monolayer. In ischemic stroke patients, calcium channel blockers improved proteinuria, suggesting that inhibition of calcium influx preserves podocyte morphology and permselectivity. Overall our results establish that TRPC6 is a novel target of HIF1α/ZEB2 axis and that transduces stroke-induced ischemia-hypoxia injury in podocytes (Fig 9).

**Figure 9.**
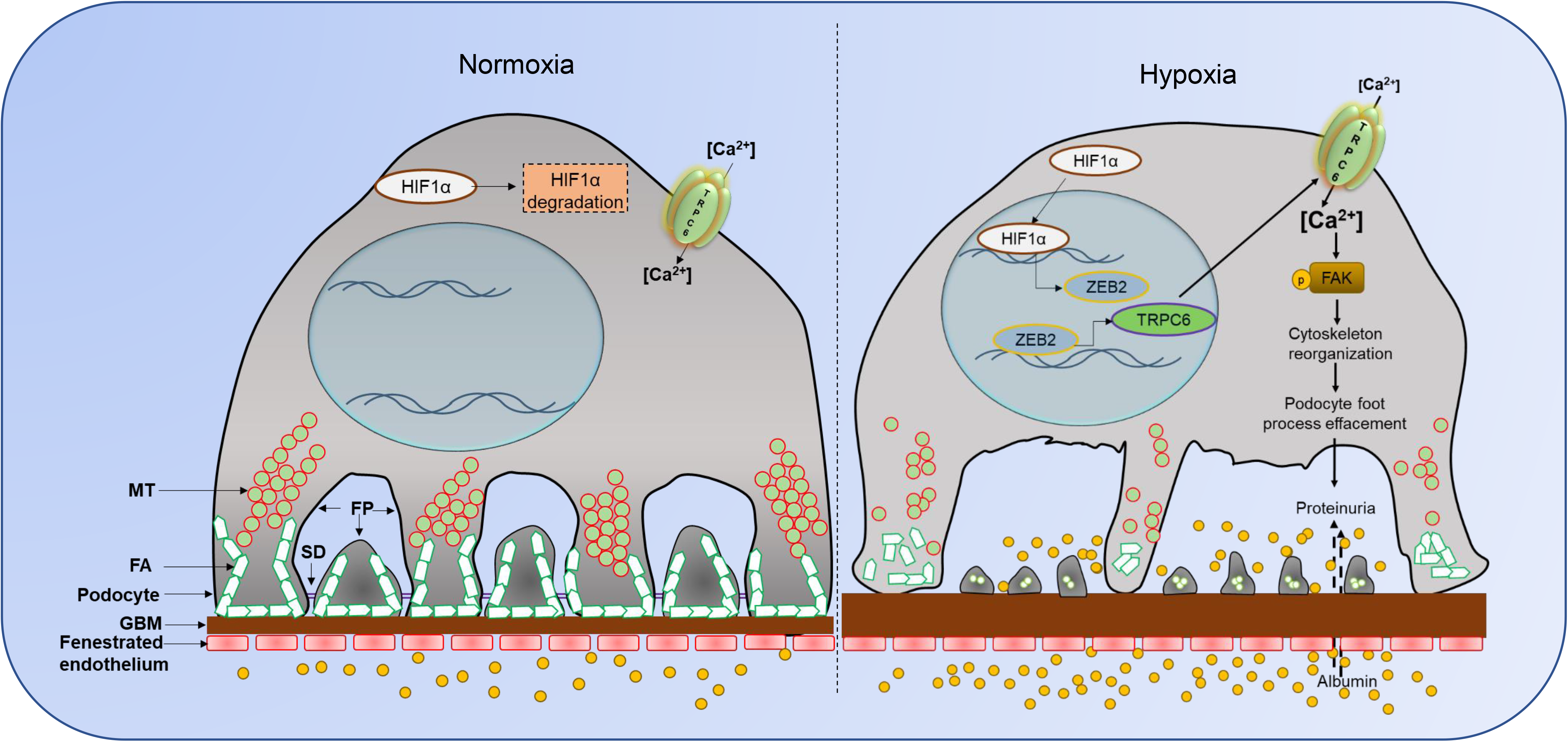
Proposed model for ischemic-hypoxia mediated podocyte injury. Ischemia-stroke rats develop systemic hypoxia that induces HIF1α accumulation in several susceptible sites including glomerular podocytes. HIF1α drives ZEB2 expression, which in turn induces TRPC6 expression. Elevated TRPC6 increases intracellular calcium levels and calcium-dependent phosphorylation of FAK elicits cytoskeletal rearrangements. These cytoskeletal rearrangements eventually manifest in effacement of podocyte foot-processes and increased permeability to proteins and large molecules. The overactivity of HIF1α-ZEB2-TRPC6 axis in podocytes elicits cytoskeletal abnormalities and proteinuria.

Rats underwent MCAO developed hypoxia as evidenced by the reduced partial pressure of oxygen (PaO2 ≤60%) and decreased oxygen saturation (SaO2≤80%) of arterial blood from 6 to 24 hours after reperfusion suggesting that these animals develop systemic hypoxia^37^. Average SaO2 levels were significantly lower in MCAO rats between 6 and 24hrs after reperfusion. MCAO is the most frequently used experimental model to mimic ischemic stroke and insufficient cerebral blood flow during ischemic stroke elicits hypoxic injury, which results in reduced arterial oxygen saturation^22^. We were interested in understanding the distant organ effect of stroke, particularly on glomerular function. Normally, synergy among arteriovenous oxygen shunting, renal blood flow, and glomerular filtration rate helps kidneys maintain arterial oxygen pressure at relatively stable levels^2^. This intricate interplay among several physiological factors makes kidneys susceptible to hypoxic injury^2^. It was reported that proteinuria is one of the major clinical outcomes following acute ischemic stroke^35^. Proteinuria refers to the impaired function of GFA; therefore, we investigated the effect of hypoxia in the glomeruli and in podocytes that are crucial to ensure glomerular permselectivity. Systemic mild hypotension is a feature of the MCAO model and mild hypotension does not affect systemic oxygen delivery to tissues^38,39^. Therefore, it appears that hypoxic effects in the MCAO group appear independent of hypotension. Although, hypertension is an independent risk factor for the stroke and it also adversely affects glomerular filtration, the management of blood pressure during the stroke period has been controversial and remains uncertain^40^. While we can’t completely rule out the role of hypertension in GFA function, in this study, we specifically focused on the effect of hypoxic signaling on glomerular podocytes.

HIF1α is a major transcription factor that transduces an array of cellular processes to let the cells adapt to hypoxic injury. Our results establish that ischemic stroke-mediated stabilization of HIF1α induces the expression of ZEB2. It was also shown earlier that ZEB2 expression is elevated in podocytes exposed to normobaric hypoxia^41^. ZEB2 related to the δEF1 protein family and these proteins possess a homeodomain flanked by N- and C-terminal zinc finger clusters^42^. Canonically, ZEB2 is considered as a transcriptional repressor. E-cadherin is a well-known target for transcriptional suppression by ZEB2. Reduced E-cadherin expression is implicated with the transition of podocytes from epithelial to mesenchymal phenotype compromising their ability to provide epithelial coverage to glomerular capillaries^23,24^. However, in this study, we found that ZEB2 induces TRPC6 expression in podocytes. Similarly, in a recent study, it was shown that ZEB2 could also serve as a transcriptional activator^27,43^. It was reported that the upstream promoter of TRPC6 has several ZEB2 binding sites^27^. Together the data suggest that ZEB2 could be a dual transcription factor with both repressor and activator functions.

Based on their involvement in the pathology of several diseases TRP channels are considered as drug targets^44^. Human TRPC sub-family of channels are most closely related to TRP channels from Drosophila. These proteins have a TRP box motif containing the consensus EWKFAR sequence at the C-terminus. These channels are in general permeable to cations whereas, TRPC6 has more affinity for calcium and excess calcium influx is implicated several pathologies. Proline to glutamine substitution at position 112 (P112Q) enhances TRPC6-mediated calcium signals in response to agonist such as angiotensin II^45^. This glutamine substitution in TRPC6 is associated with focal segmental glomerulosclerosis. Furthermore, calcium entry into cells by TRPC6 has been implicated in late-onset of Alzheimer’s disease^46^. TRPC6 is expressed in the podocyte cell body and foot processes and it interacts with slit-diaphragm proteins^47^. TRPC6 is main calcium-permeable ion channel in non-excitable cells. TRPC6 mediated calcium entry elicits albumin overload-induced ER stress and apoptosis in podocytes^48^. Other than TRP channels, cells express voltage-gated and ligand-gated calcium channels. Long-lasting type calcium channel (LTCC) are voltage-gated are classically known as dihydropyridine channel, because of the presence of a dihydropyridine-binding site. Though it is not known whether dihydropyridines inhibits TRPC6, treatment regimen with dihydropyridines improved proteinuria in stroke patients. In a recent study, it was emphasized that new dihydropyridinic calcium channel blockers can inhibit other types of calcium channels besides LTCC^44^. Furthermore, accumulated evidence suggests that dihydropyridines have pleiotropic effects to offer nephroprotection including suppressing proliferation of mesangium, attenuating mesangial entrapment of macromolecules, and countervailing the effect of platelet-derived factors^44^.

The crucial role of the cytoskeleton and structural proteins in proteinuric kidney diseases has been emphasized in several studies and notably, podocyte actin cytoskeletal rearrangement is the common final pathway subsequently leads to podocyte foot process effacement^49^. The contractile actin filament bundles that are highly ordered and arranged parallel in the foot processes were converted into disordered, short, and branched under pathological conditions, thus ensuring podocytes to compromise their unique structure. The disruption of stress fibers of the podocyte actin cytoskeleton could probably explain the reason for altered cell morphology and FPE in podocytes from ischemic reperfusion injury. Disordered actin cytoskeleton was also observed in some transient proteinuric models such as the protamine sulfate infusion and lipopolysaccharide injection, where reversible FPE and proteinuria was observed^50,51^. FAK is a non-receptor tyrosine kinase, which is recruited to focal adhesions by paxillin and talin^52^. FAK plays an essential role in cell motility, maintenance of cell morphology, and also regulates podocyte cytoskeleton. Podocyte-specific deletion of FAK in mice protected from podocyte injury and proteinuria^30^. A recent study suggests that podocyte injury activates FAK phosphorylation that elicits increased FAK turnover, FPE, and proteinuria^30^. FAK phosphorylation was observed during podocyte injury and it suggests that inhibition of FAK signaling cascade may have therapeutic potential in the treatment of glomerular injury^30^.

Previous studies have been shown that acute ischemic stroke induces intracellular calcium accumulation^53^. The increased overload of intracellular calcium in stroke results in cell death by necrosis and cell catabolism^54^. Administration of calcium channel blockers improved renal function, GFR, renal blood flow, and electrolyte excretion^55^. Other reports have shown that CCBs are involved in exerting the positive hemodynamic function in diabetes and clinical trials demonstrated reduced urinary protein excretion^56^. Ischemic stroke with hypertension together with proteinuria is one of the predominant factors in contributing to the progression of CKD, which is a significant determinant of mortality and morbidity among stroke patients^57^. Stroke patients treated with calcium channel blockers appear to show reduced proteinuria and UACR. In summary, our study identified TRPC6 as a bona fide target of ZEB2 and transduces ischemia mediated podocyte injury and proteinuria. TRPC6 mediated calcium influx possibly mediate the podocyte cytoskeletal abnormality and calcium blockers could be a therapeutic option to combat ischemia-hypoxia injury in podocytes.

## Materials and Methods

### Intraluminal suture middle cerebral artery occlusion

Middle cerebral artery occlusion (MCAO) and sham surgery were performed in 8-week-old Male SD rats as reported earlier using the intraluminal monofilament technique^58^. Cerebral reperfusion was allowed by withdrawing the monofilament carefully after 2h of surgery and animals were maintained for 24h. The rats subjected to stroke and sham-surgery were transcardially perfused with saline under anesthetic conditions. Tissues were harvested and frozen for RNA and protein isolation whereas for fresh tissues were fixed for histological studies and TEM imaging. To assess the success of the model and infarct volume, 2-mm-thick coronal sections of the brain were prepared and stained with 1% triphenyltetrazolium chloride. University of Hyderabad’s Institutional animal ethics committee approved animal experiment protocols. We confirm that all methods were performed in accordance with the relevant guidelines and regulations.

### Urine analysis

Urine samples from rats were subjected to 10% SDS-PAGE gel and processed for silver staining as described earlier^23^. Urinary albumin (#11573) and creatinine (#11502) were measured using commercially available kits (Biosystems, Barcelona, Spain).

### Isolation of glomeruli and podocytes

The glomeruli from the kidneys were isolated by a series of stainless sieves as described earlier^23^. Primary podocytes from rat kidney were isolated as reported earlier^59^.

### Immunohistochemistry

Paraffin sections (5μm) of kidney cortex were prepared with Leica microtome on to pre-coated glass slides. Sections were allowed for deparaffinization, rehydration, followed by antigen retrieval. Following permeabilization and blocking, sections were incubated with respective primary antibody at 4°C for overnight. Further, incubation with secondary antibody and DAB staining with the kit method using Mouse/Rabbit PolyDetector DAB-HRP Detection kit (Santa Barbara, CA, USA). Images were captured with Trinocular microscope 100X objective (Leica, Buffalo Grove, IL).

### Transmission Electron microscopy

Kidney sections were processed for TEM imaging as described in our earlier study^23^. Briefly, the renal cortex portion of the kidney from sham and stroke-induced rats were collected and fixed with 2.5% glutaraldehyde and 1% osmium tetroxide. Ultrathin sections were mounted on copper grids, stained with 3% uranyl acetate and images were obtained with JEM-1400TEM (Jeol, Peabody, MA).

### Cell Culture

In this study, we employed human podocytes (A gift from Prof. Moin Saleem, University of Bristol) and HEK293T cells. Podocytes and HEK293T cells were cultured essentially as described earlier^23,24^.

### Immunoblotting

Equal concentration of protein from either glomerular or cell lysate was subjected to SDS-PAGE and blotted onto the nitrocellulose membrane. Immunoblotting and developing the blots was performed as reported earlier^23^.

### qRT-PCR analysis

Isolation of RNA, preparation of cDNA, and qRT-PCR were performed as reported earlier^41^. The expression level of each mRNA was normalized to β-Actin and quantified using the comparative Ct method. List of primers used in this study is provided in Table S3.

### Immunofluorescence

Podocytes were cultured on coverslips and allowed to differentiate. Following experimental conditions, these cells fixed with 4% paraformaldehyde and performed immunofluorescence protocol as reported earlier^23^. Imaging was done in a Leica trinocular fluorescent microscope under 60x oil objective.

### Transfection

HEK293T cells or differentiated podocytes were transfected using jetPEI reagent (Polyplus, Illkirch, France). 1×10^5^ cells were seeded per well in 6-well cell culture plates and transiently transfected with plasmid DNA or siRNA by mixing with NaCl-jetPEI complexes. After 48 hr of transfection, cells were washed twice with PBS and lysed with RIPA buffer; the expression levels were measured by western blotting as described above. Podocytes that ectopically express ZEB2 or ZEB2 knockdown were employed in calcium influx assay.

### Measurement of Intracellular Calcium

Fluo3-AM (Sigma#92210) is a highly sensitive cell permeable dye that binds selectively to calcium^60^. Fluo3-AM is non-fluorescent until it is hydrolyzed intracellularly and/or in the presence of calcium. Differentiated podocytes were treated with FG-4592 in the presence or absence of calcium channel blocker (2APB) for 4h and were loaded with 0.5mM Fluo3-AM and incubated for 1hr at 37ºC with 5%CO2. Cells were then washed with calcium-free PBS three times to remove the extracellular Fluo3-AM. Cells were lysed with 1% NP-40 and 0.5M EGTA and fluorescence of intracellular calcium bound Fluo3-AM was recorded using fluorometer (λex 506 nm; λem 525 nm). Intracellular calcium was also measured in podocytes that overexpress ZEB2 or in podocytes in which ZEB2 was knockdown in the presence or absence of FG-4592 and 2APB.

### Phalloidin staining

Fluorescent phalloidin-TRITC conjugate staining was performed to visualize the distribution of stress fibers in differentiated podocytes essentially as described earlier ^41^.

### ChIP Assay

Approximately 80% of confluent podocytes were exposed to hypoxia. Chromatin immunoprecipitation (ChIP) was performed as described earlier^23^. Cells were cross-linked with formaldehyde and quenched with 125mM glycine, then lysed with lysis buffer (1% SDS, 10mMEDTA, 50mM Tris-HCl, pH-8.1, protease inhibitors). Pre-clearing was performed with protein A/G beads followed by immunoprecipitation with the antibody of interest. Chromatin fragments were eluted from the beads and further purified by the phenol-chloroform method. This purified DNA was used for RT-PCR to quantify fold enrichment. The list of primers used in for ChiP experiments is provided in Table S4.

### Promoter-Reporter Assay

TRPC6 promoter was cloned into PGL3 basic reporter vector with MluI and BglII restriction sites. The resultant pGL3-TRPC6 promoter construct was co-transfected with renilla plasmid into the HEK293T cells. Cells were then exposed to hypoxia or transfected with ZEB2 are scrapped with ice-cold PBS. Cells were collected after a spin down at 5,000 rpm for 5min at 4°C, followed by lysing the cells with a passive lysis buffer. 20ul of lysate was used for measuring luciferase expression with LARII and renilla expression with stop glow reagents to quantify relative promoter activity.

### Albumin influx assay

Podocyte permeability was assessed by albumin influx assay as described earlier^23^. Briefly, human podocytes were allowed to differentiate as described above on collagen-coated transwell filters. Differentiated podocytes were exposed to hypoxia for 24 hrs and washed with phosphate buffer containing with 1mM MgCl_2_ and 1mM CaCl_2_. In the bottom chamber, 2 ml of RMPI 1640 medium containing 40 mg/ml BSA was added. In the top chamber, only RPMI 1640 (without albumin) was added. The concentration of BSA in the upper chamber was estimated at various time points.

### Human Patients Data

The ischemic stroke study protocol was approved by the institutional review board of Narayana medical college (Nellore, India). The study was conducted in accordance with the declaration of Helsinki guidelines. Patients provided written informed consent during enrolment in the study. A total of 120 Ischemic stroke patients (Table S1) were recruited in this study and those patients who had other inference, such as CVDs, diabetes, and hypertension were excluded from the study. The stroke patients were from southern coastal Andhra Pradesh, India. Urine samples and kidney function parameters were measured in stroke patients before and after treatment with CCBs. Mean treatment regimen follow-up period was 3yrs (Table S2).

### Statistical analysis

Statistical analyses were performed in GraphPad Prism V.7. For all the statistical differences, we used a 2-tailed, unpaired Student’s t-test or Mann-Whitney U test to analyze the differences between 2 groups. The Bonferroni correction was used when more than >2 groups were present. Data are presented as mean ± S.E. unless otherwise indicated. p values equal to or less than 0.05 were considered significant.

## Acknowledgments

Life Science Research Board (LSRB-296) of DRDO, INDIA and Indian Council of Medical Research (2019-0905) to AKP funded this study. Fellowship by Indian Council of Medical Research (to KMN) and University Grants Commission (to RN, DM, and LPK) are acknowledged.

## Author contributions

KN, RN, and AKP conceived and designed research; KN, RN, DM, SKT, VPN, and LPK performed experiments; KN, RN, DM, and AKP analyzed the data; PBP, PN, and SSKN obtained human patient’s data; KN, RN, SSKN, PBP, and AKP interpreted the data; KN, RN, and AKP wrote the manuscript with the inputs from the rest of the authors.

