## Supplementary data for "Cerebral ischemia induces TRPC6 in the glomerular podocytes: A novel role for HIF1α/ZEB2 axis in the pathogenesis of stroke-induced proteinuria"

**Figure 9.** Proposed model for ischemic-hypoxia mediated podocyte injury. Ischemia-stroke rats develop systemic hypoxia that induces HIF1α accumulation in several susceptible sites including glomerular podocytes. HIF1α drives ZEB2 expression, which in turn induces TRPC6 expression. Elevated TRPC6 increases intracellular calcium levels and calcium-dependent phosphorylation of FAK elicits cytoskeletal rearrangements. These cytoskeletal rearrangements eventually manifest in effacement of podocyte foot-processes and increased permeability to proteins and large molecules. The overactivity of HIF1α-ZEB2-TRPC6 axis in podocytes elicits cytoskeletal abnormalities and proteinuria.

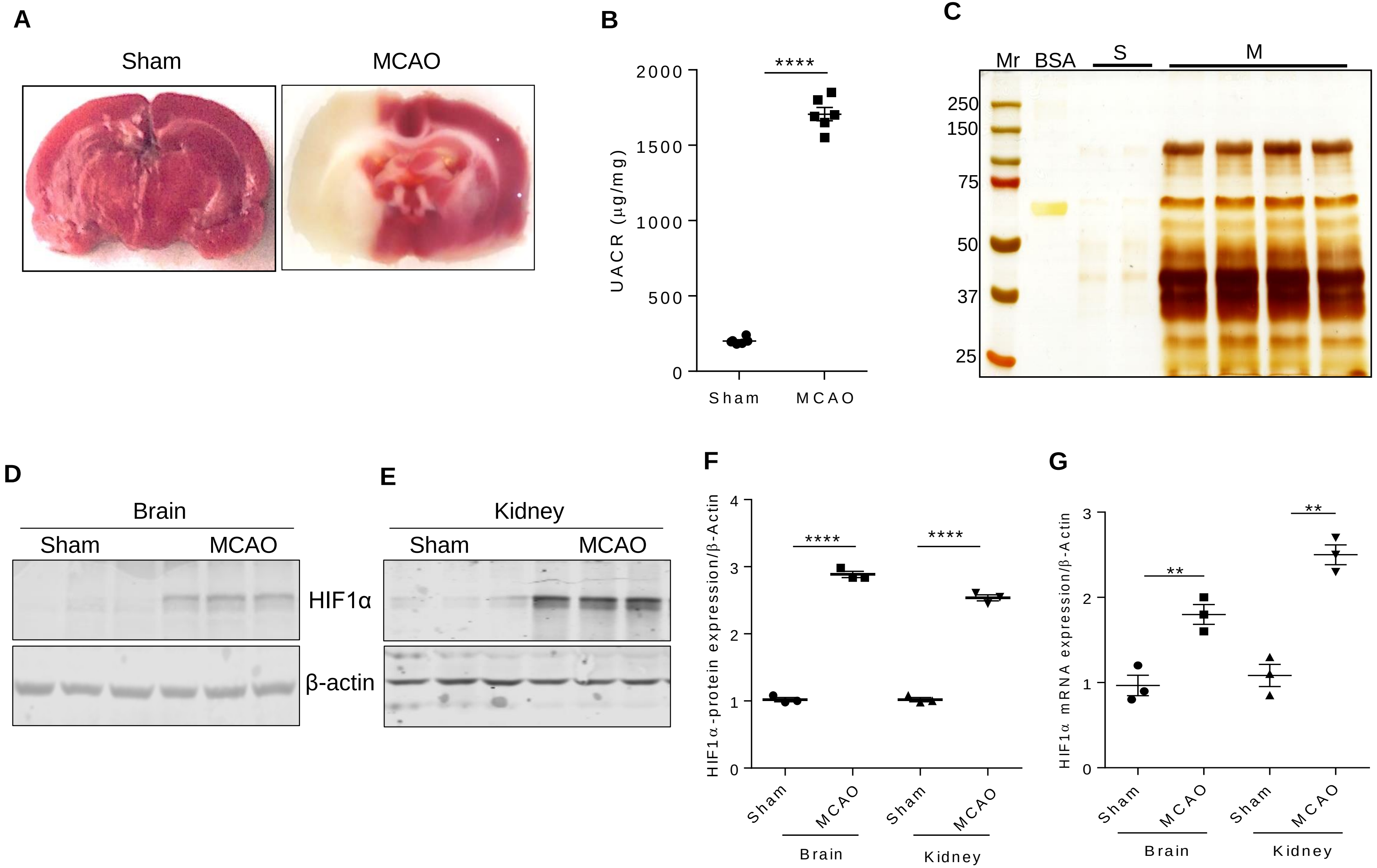
A
Sham
MCAO
B
****
2000
1500
1000
500
0
UACR (μg/mg)
Sham
MCAO
C
Mr
BSA
S
M
250
150
75
50
37
25
D
Brain
Sham
MCAO
HIF1α
β-actin
E
Kidney
Sham
MCAO
F
HIF1α-protein expression/β-Actin
4
3
2
1
0
****
****
Sham
MCAO
Sham
MCAO
Brain
Kidney
G
HIF1α mRNA expression/β-Actin
3
2
1
0
**
**
Sham
MCAO
Sham
MCAO
Brain
Kidney

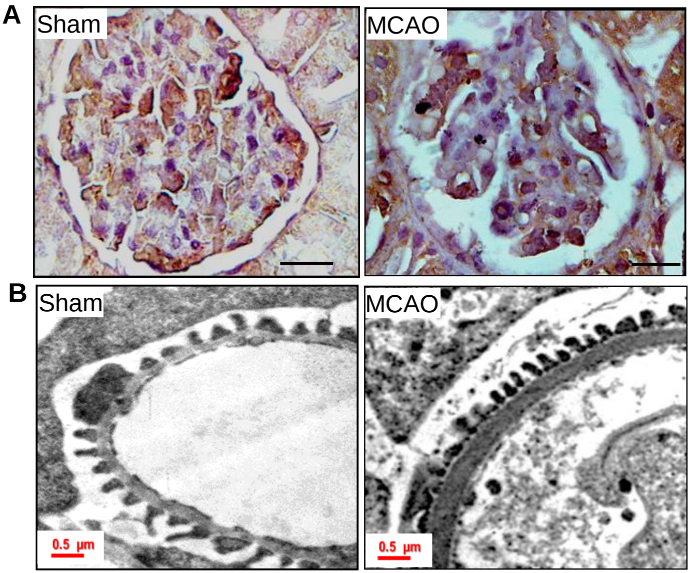

A
Sham
MCAO
B
Sham
MCAO
0.5 µm
0.5 µm

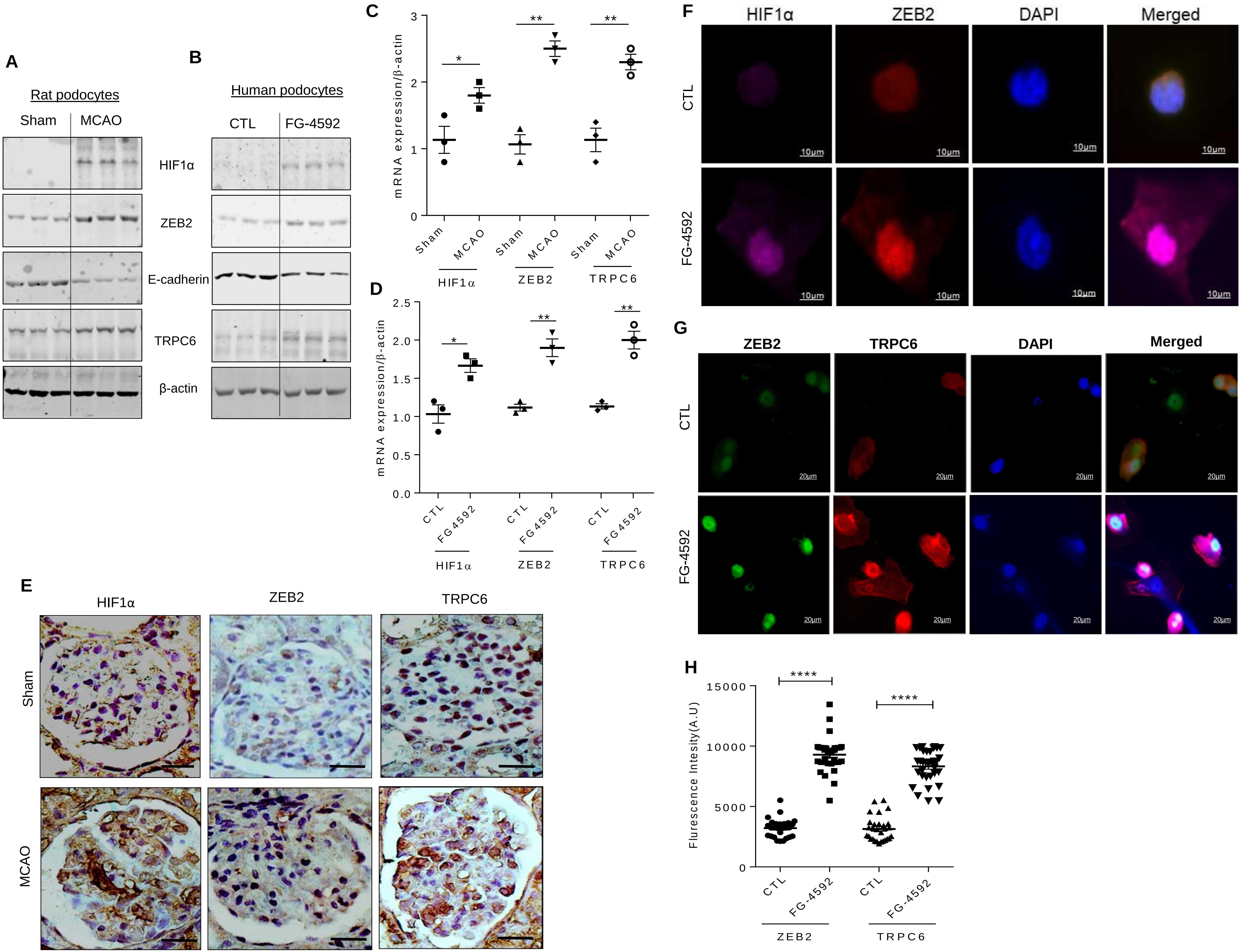
A
Rat podocytes
Sham
MCAO
HIF1α
ZEB2
E-cadherin
TRPC6
β-actin
B
Human podocytes
CTL
FG-4592
C
mRNA expression/β-actin
Sham
MCAO
HIF1α
ZEB2
TRPC6
D
mRNA expression/β-actin
CTL
FG 4592
E
HIF1α
ZEB2
TRPC6
Sham
MCAO
F
HIF1α
ZEB2
DAPI
Merged
CTL
FG-4592
10μm
G
ZEB2
TRPC6
DAPI
Merged
CTL
FG-4592
20μm
H
Flurescence Intesity(A.U)
CTL
FG-4592
ZEB2
TRPC6

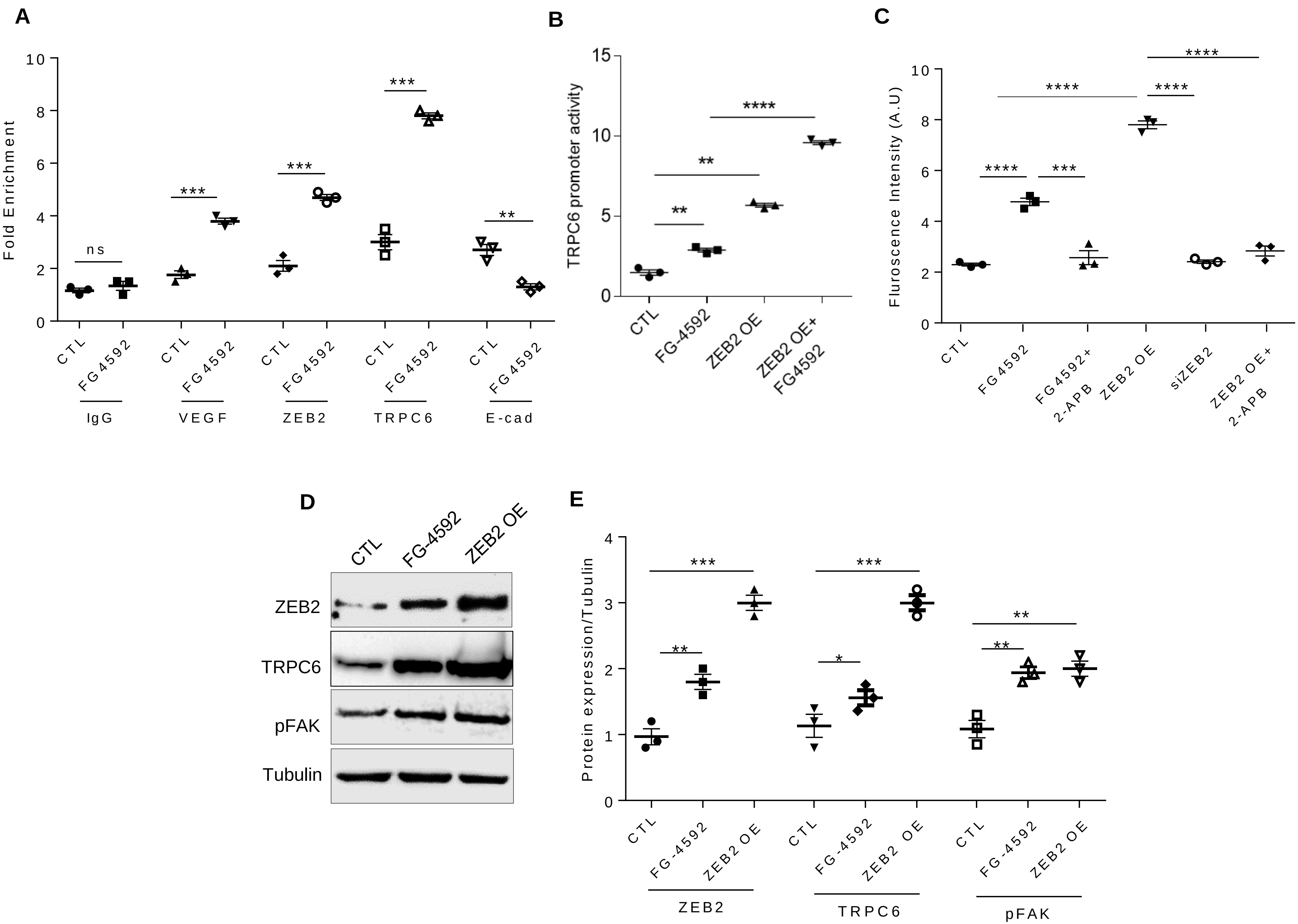

A
Fold Enrichment
ns
***
***
***
**
CTL
FG4592
IgG
VEGF
ZEB2
TRPC6
E-cad
B
TRPC6 promoter activity
**
**
****
CTL
FG-4592
ZEB2 OE
ZEB2 OE+
FG4592
C
Fluroscence Intensity (A.U)
****
***
****
****
****
CTL
FG4592
FG4592+
2-APB
ZEB2 OE
siZEB2
ZEB2 OE+
2-APB
D
CTL
FG-4592
ZEB2 OE
ZEB2
TRPC6
pFAK
Tubulin
E
Protein expression/Tubulin
***
**
***
*
**
**
CTL
FG-4592
ZEB2 OE
ZEB2
TRPC6
pFAK

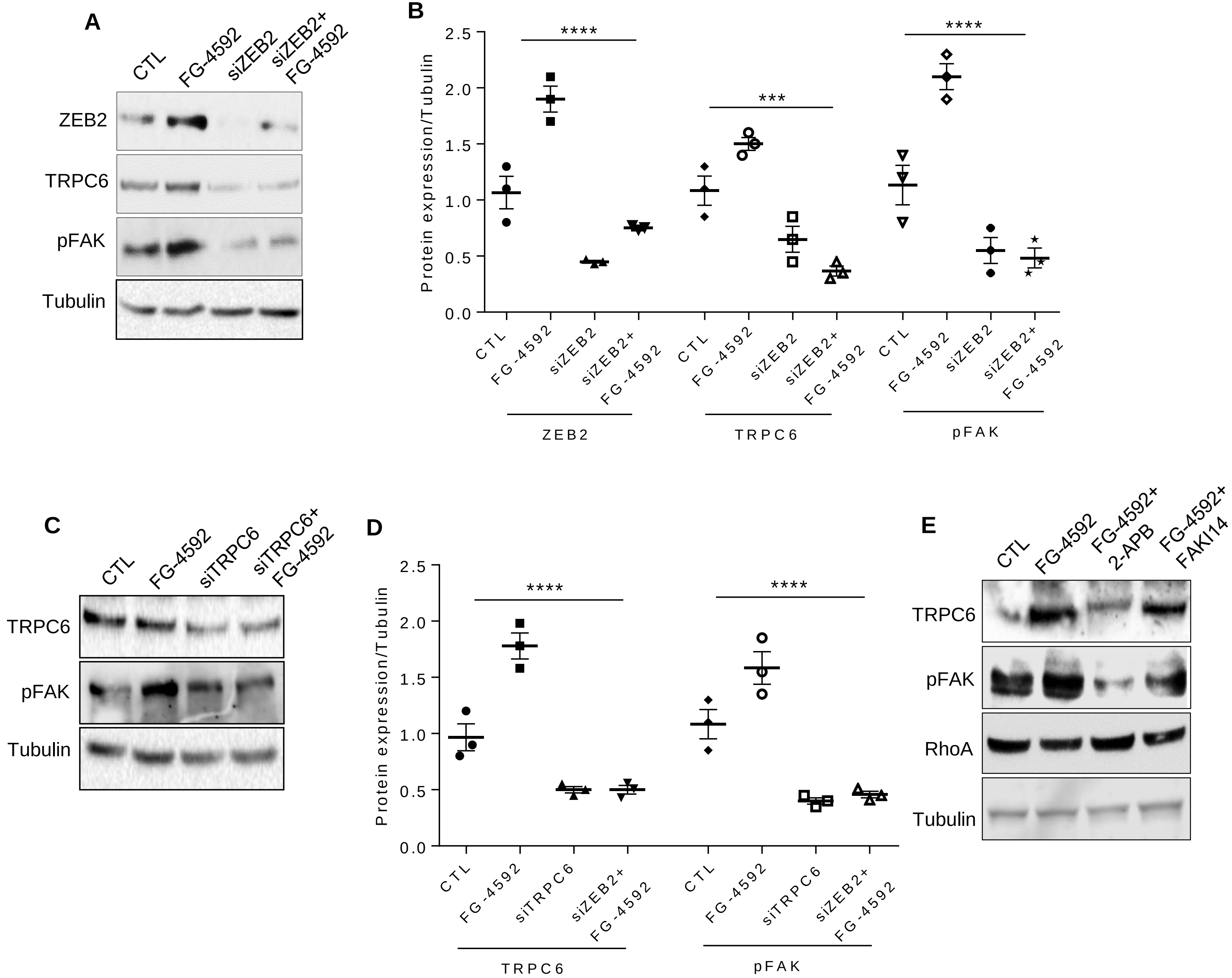
A
CTL
FG-4592
siZEB2
siZEB2+ FG-4592
ZEB2
TRPC6
pFAK
Tubulin
B
Protein expression/Tubulin
2.5
2.0
1.5
1.0
0.5
0.0
****
***
****
CTL
FG-4592
siZEB2
siZEB2+ FG-4592
ZEB2
TRPC6
pFAK
C
CTL
FG-4592
siTRPC6
siTRPC6+ FG-4592
TRPC6
pFAK
Tubulin
D
Protein expression/Tubulin
****
****
CTL
FG-4592
siTRPC6
siZEB2+ FG-4592
TRPC6
pFAK
E
CTL
FG-4592
FG-4592+ 2-APB
FG-4592+ FAKI14
TRPC6
pFAK
RhoA
Tubulin

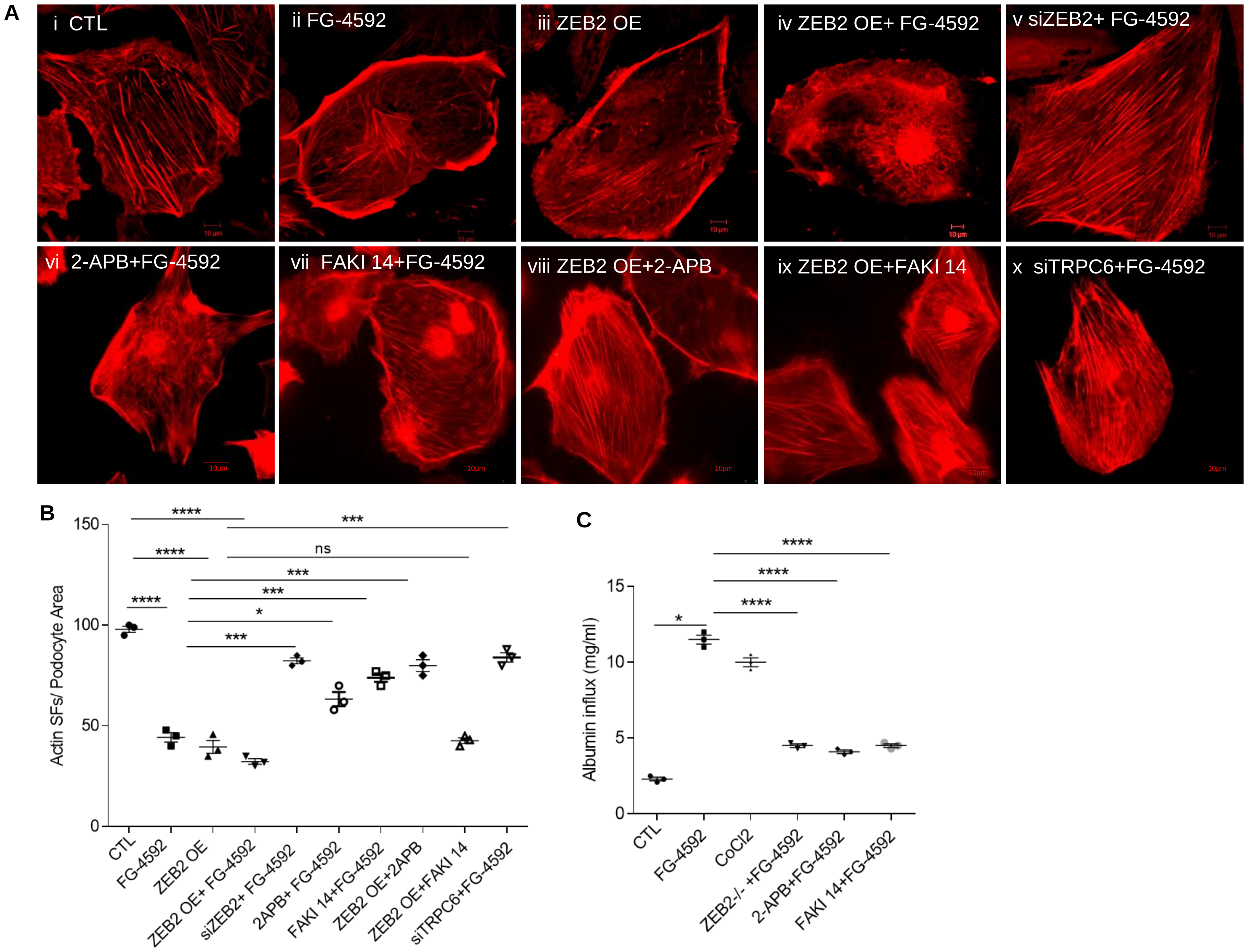
A
i CTL
ii FG-4592
iii ZEB2 OE
iv ZEB2 OE+ FG-4592
v siZEB2+ FG-4592
vi 2-APB+FG-4592
vii FAKI 14+FG-4592
viii ZEB2 OE+2-APB
ix ZEB2 OE+FAKI 14
x siTRPC6+FG-4592
B
Actin SFs/ Podocyte Area
0
50
100
150
****
****
****
***
ns
***
***
*
***
CTL
FG-4592
ZEB2 OE
ZEB2 OE+ FG-4592
siZEB2+ FG-4592
2APB+ FG-4592
FAKI 14+FG-4592
ZEB2 OE+2APB
ZEB2 OE+FAKI 14
siTRPC6+FG-4592
C
Albumin influx (mg/ml)
0
5
10
15
*
****
****
****
CTL
FG-4592
CoCl2
ZEB2-/- +FG-4592
2-APB+FG-4592
FAKI 14+FG-4592

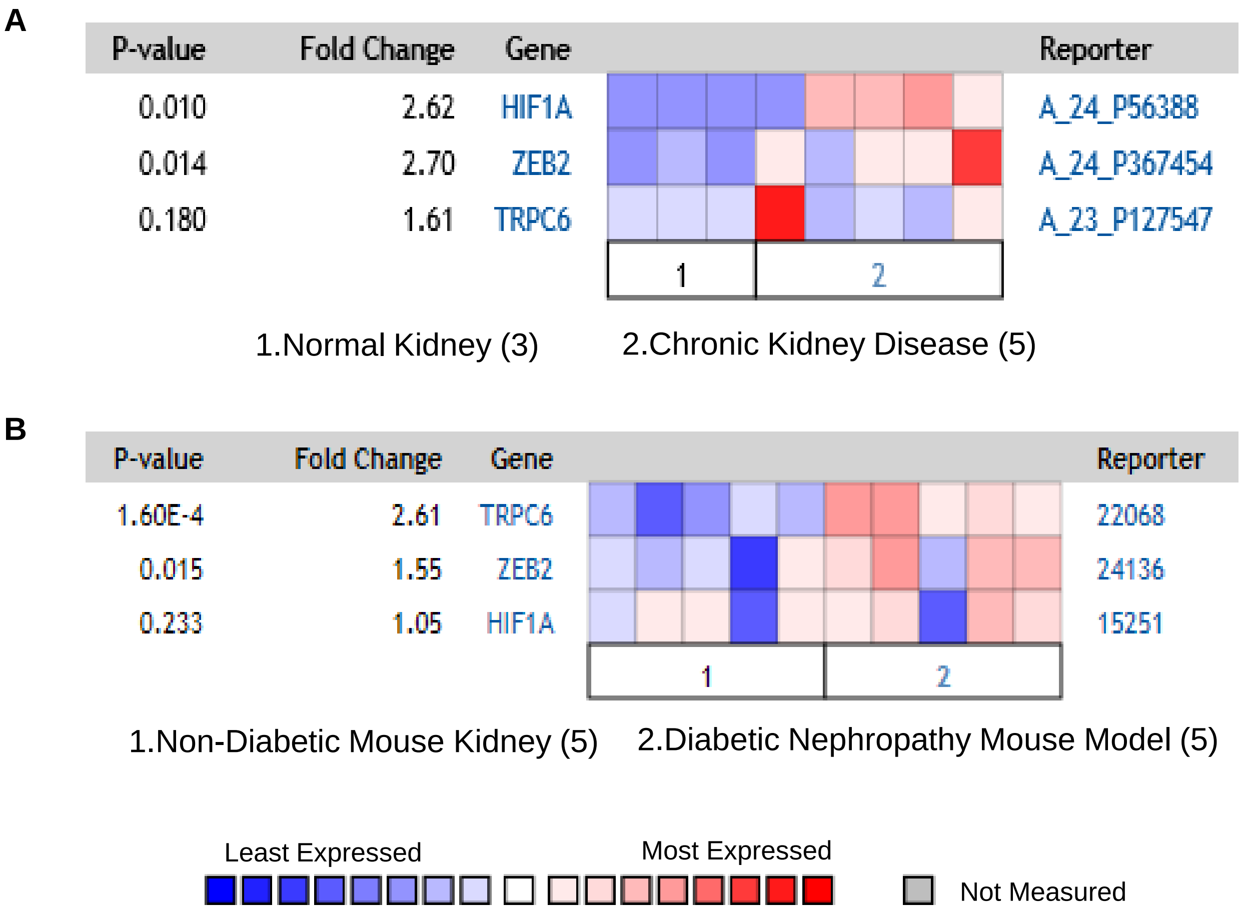
A
P-value
Fold Change
Gene
Reporter
0.010
2.62
HIF1A
A_24_P56388
0.014
2.70
ZEB2
A_24_P367454
0.180
1.61
TRPC6
A_23_P127547
1
2
1.Normal Kidney (3)
2.Chronic Kidney Disease (5)
B
P-value
Fold Change
Gene
Reporter
1.60E-4
2.61
TRPC6
22068
0.015
1.55
ZEB2
24136
0.233
1.05
HIF1A
15251
1
2
1.Non-Diabetic Mouse Kidney (5)
2.Diabetic Nephropathy Mouse Model (5)
Least Expressed
Most Expressed
Not Measured

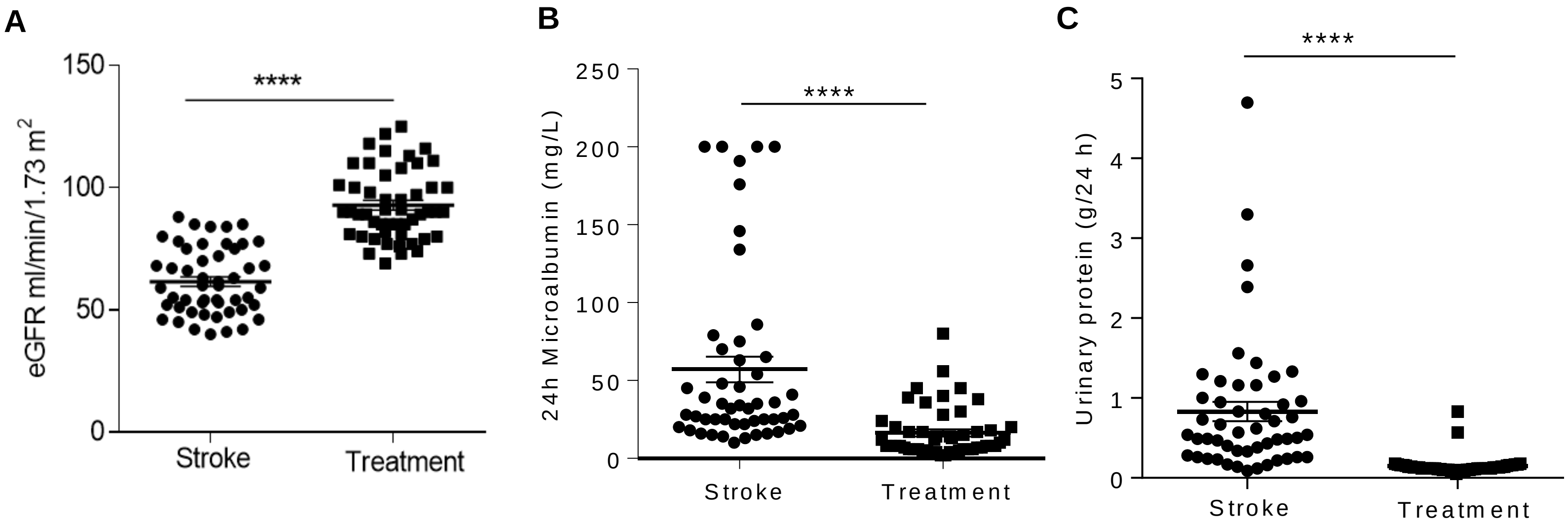
A
eGFR ml/min/1.73 m2
150
100
50
0
****
Stroke
Treatment
B
24h Microalbumin (mg/L)
250
200
150
100
50
0
****
Stroke
Treatment
C
Urinary protein (g/24 h)
5
4
3
2
1
0
****
Stroke
Treatment

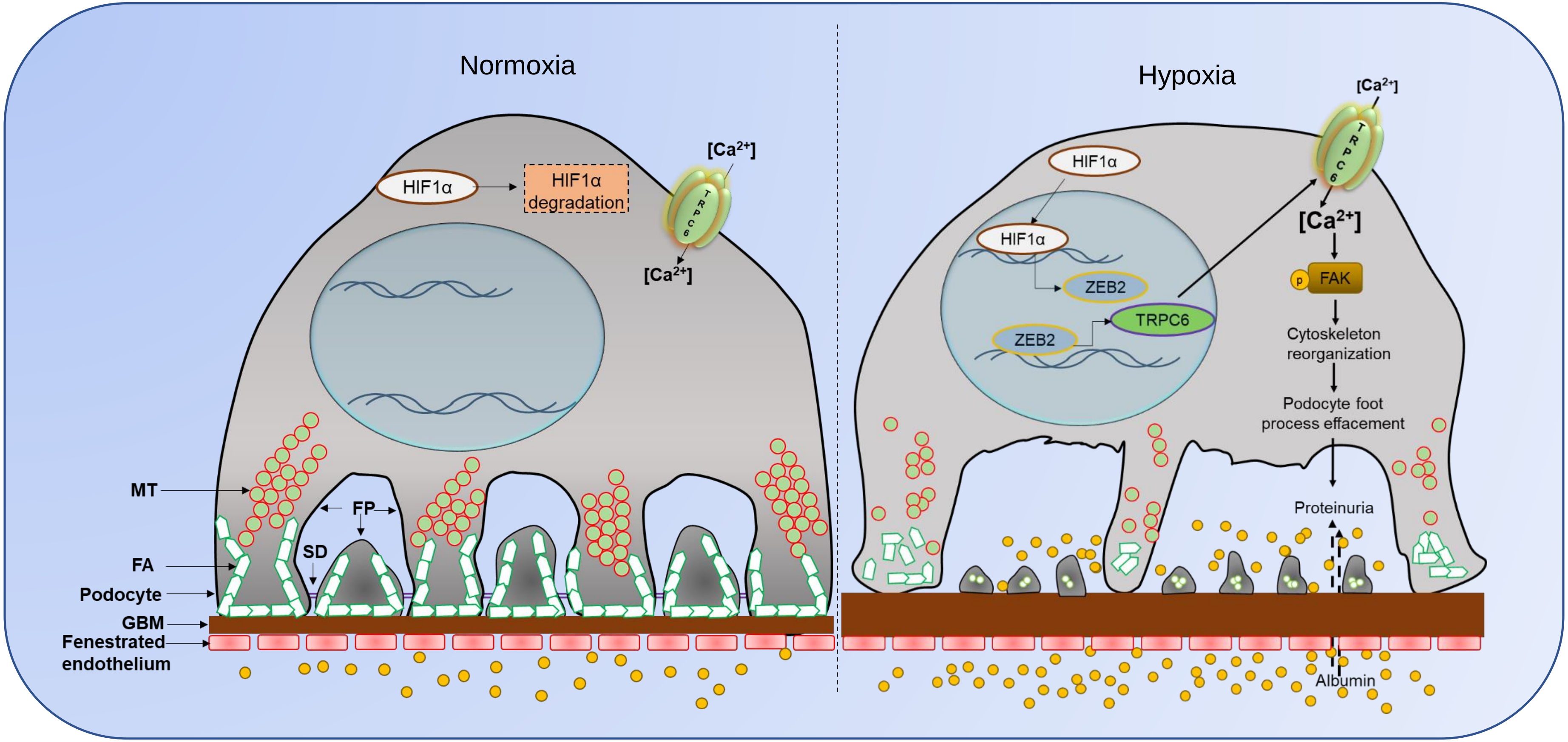
Normoxia
Hypoxia
HIF1α
HIF1α degradation
[Ca2+]
TRPC6
[Ca2+]
MT
FP
SD
FA
Podocyte
GBM
Fenestrated endothelium
[Ca2+]
HIF1α
HIF1α
ZEB2
ZEB2
TRPC6
[Ca2+]
p
FAK
Cytoskeleton reorganization
Podocyte foot process effacement
Proteinuria
Albumin
